# Regulation of coronavirus nsp15 cleavage specificity by RNA structure

**DOI:** 10.1101/2023.05.12.540483

**Authors:** Indranil Salukhe, Ryan Choi, Wesley Van Voorhis, Lynn Barrett, Jennifer L. Hyde

## Abstract

SARS-CoV-2, the etiologic agent of the COVID-19 pandemic, has had an enduring impact on global public health. However, SARS-CoV-2 is only one of multiple pathogenic human coronaviruses (CoVs) to have emerged since the turn of the century. CoVs encode for several nonstructural proteins (NSPS) that are essential for viral replication and pathogenesis. Among them is nsp15, a uridine-specific viral endonuclease that is important in evading the host immune response and promoting viral replication. Despite the established function of nsp15 as a uridine-specific endonuclease, little is known about other determinants of its cleavage specificity. In this study we investigate the role of RNA secondary structure in SARS-CoV-2 nsp15 endonuclease activity.

Using a series of *in vitro* endonuclease assays, we observed that thermodynamically stable RNA structures were protected from nsp15 cleavage relative to RNAs lacking stable structure. We leveraged the s2m RNA from the SARS 3’UTR as a model for our structural studies as it adopts a well-defined structure with several uridines, two of which are unpaired and thus high probably targets for nsp15 cleavage. We found that SARS-CoV-2 nsp15 specifically cleaves s2m at the unpaired uridine within the GNRNA pentaloop of the RNA. Further investigation revealed that the position of uridine within the pentaloop also impacted nsp15 cleavage efficiency, suggesting that positioning within the pentaloop is necessary for optimal presentation of the scissile uridine and alignment within the nsp15 catalytic pocket. Our findings indicate that RNA secondary structure is an important determinant of nsp15 cleavage and provides insight into the molecular mechanisms of recognition of RNA by nsp15.

## Introduction

Coronaviruses (CoVs) are a diverse group of enveloped, positive sense single-stranded RNA (+ssRNA) viruses within the *Nidovirales* order which infect a wide range of host species and are associated with mild to severe disease in livestock and humans (1–3). The *Coronaviridae* family is composed of four genera: Alpha-, Beta-, Gamma-, and Deltacoronaviruses (4). Since the turn of the century, several pathogenic CoVs have emerged in the human population including SARS, MERS, and SARS-CoV-2 with the latter being the causative agent of the ongoing COVID-19 pandemic (1, 2).

Replication of the large (∼30kb) CoV genome is facilitated by multiple nonstructural proteins (nsp1-16) which are produced from two open reading frames encoded by the first two thirds of the viral genome. These proteins encode several key functions necessary for synthesis and post-translational processing of the viral genome, including RNA-dependent RNA polymerase (nsp12), helicase (nsp13), RNA M7 and 2D-O-methyltransferase (nsp14, nsp16/nsp10), protease (nsp3, nsp5), and exonuclease (nsp14) activities. Structural proteins which assemble to form the virion, as well as accessory proteins, are encoded in a set of nested subgenomic RNAs which are coterminal with the 3’ end of the genome.

The viral replication/transcription complex (RTC) is housed within membrane structures derived from the host endoplasmic reticulum (ER) which are induced by expression of nsp3, 4, and 6 (5–12). Like other +ssRNA viruses, these double-membrane vesicles (DMVs) provide a microenvironment that supports efficient viral replication and sequester double-stranded RNA (dsRNA) replication intermediates from detection by host pathogen recognition receptors (PRRs) (13, 14). Previous studies have demonstrated a key role for the interferon (IFN) response in restriction of CoVs. Accordingly, CoVs have evolved several mechanisms to evade antiviral immunity (e.g., 2D-O-methylation) and encode several proteins which antagonize the IFN response (nsp1, ORF6) (15). Nsp15 is a uridine-specific endonuclease (endoU) encoded by CoVs and other Nidoviruses which has been implicated in evasion of host-mediated RNA sensing and IFN activation (16, 17). While nsp15 has been shown to be non-essential for viral replication of several CoVs, absence of nsp15 endoU activity (Δnsp15) leads to greater accumulation of dsRNA replication intermediates which trigger IFN activation in immune competent cells such as macrophages (16). During CoV infection, dsRNA sensing occurs primarily via MDA-5, though other RNA sensors (PKR) have also been implicated (16, 18).

While originally described as a uridine-specific endonuclease, nsp15 has been shown to preferentially cleave pyrimidine-adenine dinucleotides (U^↓^A or C^↓^A) (19). More recently, viral targets of nsp15 have been identified in both the polyU tract of tract of 5’ negative-sense RNA, as well as at several sites throughout the CoV genome (17, 19). Although multiple nsp15 cleavage sites have been mapped throughout the viral genome, no other sequence determinants have yet been identified. Despite the abundance of pyrimidine-adenine dinucleotides across the SARS-CoV-2 genome, only a fraction of these sites are cleaved by nsp15, suggesting that other RNA determinants likely affect cleavage specificity of viral RNA. Earlier *in vitro* studies have shown that SARS nsp15 cleaves a single unpaired uridine in the stem-loop II motif (s2m) RNA structural element found in the 3’-UTR of the viral genome suggesting that the structural context in which the scissile uridine is presented may play a role in determining nsp15 cleavage specificity (20). In this study, we use SARS s2m as a model RNA to explore the secondary structure requirements for SARS-CoV-2 nsp15 cleavage. Our findings reveal that thermodynamically stable RNAs are protected from SARS-CoV-2 nsp15 cleavage and cleaved less efficiently. We show for the first time that SARS-CoV-2 nsp15 preferentially cleaves unpaired bases. and show that the positioning and sequence context of the scissile uridine play a role in determining SARS-CoV-2 nsp15 cleavage specificity. This is consistent with previous studies of SARS nsp15, suggesting that SARS-CoV-2, and CoVs more broadly, may use RNA structure to regulate nsp15 cleavage specificity. These data shed further light on requirements for viral endoU target recognition and demonstrate an underappreciated role for viral RNA structure in evasion of innate immunity.

## Results

### RNA secondary structure modulates nsp15 cleavage efficiency

Nsp15 cleavage sites have previously been mapped in the MHV genome following viral infection (17, 19). Despite the abundance of pyrimidine-adenine dinucleotides throughout the genome, MHV nsp15 was found to cleave only a fraction of these sites, suggesting that other determinants play a role in CoV nsp15 cleavage specificity. We analyzed sequences upstream and downstream of MHV nsp15 cleavage sites to identify shared motifs that might contribute to nsp15 recognition and specificity. We reasoned that putative nsp15 recognition motifs present in the genome of MHV may also be conserved across other betacoronaviruses, including SARS-CoV-2. Using the MEME-suite XSTREME motif analysis and discovery tool (21) we analyzed 200 nt sequences corresponding to 100 nt upstream and downstream of the MHV nsp15 cleavage sites identified by Ancar et al. Although some motifs were found to be enriched in sequences surrounding the MHV nsp15 cleavage sites, they were not uniformly conserved across all cleavage sites. Moreover, many of these motifs were found at other sites in the viral genome which were not previously identified as nsp15 targets and were not found to be conserved in SARS-CoV-2 or other human CoVs.

As RNA structure has previously been implicated in nsp15 cleavage specificity (20), we next compared the underlying predicted secondary structure of the same 200 nt fragments using LocRNA software (22, 23). We did not identify any conserved structural elements at nsp15 cleavage sites in MHV nor did we identify conserved structural elements upstream or downstream of these sites. Despite the absence of any conserved sequence or structural motifs, we hypothesized that the presence of thermodynamically stable RNA elements would prevent cleavage of RNA by nsp15.

To test this hypothesis, we used RNAfold (24) to identify regions across the SARS-CoV-2 genome predicted to have varying degrees of thermodynamic stability (Fig. 1A; RNA 1, RNA 2, RNA 3) (25, 26). We identified representative regions of high, moderate, and low thermodynamic stability as indicated by their relative ΔG values (Fig. 1B), with more stable predicted structures having lower ΔG values (27). 100 nt RNAs corresponding to these regions were synthesized *in vitro*, and cleavage efficiency of these RNAs was assessed using a SARS-CoV-2 nsp15 endonuclease assay (28) (Fig. 1C and D). Folded RNAs were incubated with recombinant SARS-CoV-2 nsp15 for varying times (1, 15, 60, and 240 minutes) and cleavage products analyzed by polyacrylamide gel electrophoresis (Fig. 1C). Cleavage of full-length RNA was additionally quantified by densitometry by comparing the abundance of full-length RNA at each time point to a denatured nsp15 control (no cleavage, (-) far-right lane) (Fig. 1D). As we hypothesized, we observed that RNAs with relatively moderate or low predicted thermodynamic stability were cleaved more rapidly than RNA with relatively high thermodynamically stability (Fig. 1C, compare abundance of full-length transcript at 1, 15 and 60 min [red arrow]; Fig. 1D compare percentage uncleaved RNA). Notably, RNA with relatively low thermodynamic stability showed rapid cleavage as early as 1 min following the addition of nsp15. While each RNA yielded different cleavage products owing to the differences in RNA sequence, we observed the appearance of distinct cleavage products that either decreased (green arrows) or increased (blue arrows) in abundance over time. These likely represent cleavage intermediates or end products respectively, suggesting that other structural features or specific sequences also likely contribute to the higher efficiency of nsp15 cleavage of some RNA sequences over others. Collectively, these data suggest that RNA secondary structure impacts nsp15-dependent RNA cleavage.

**Figure 1.**
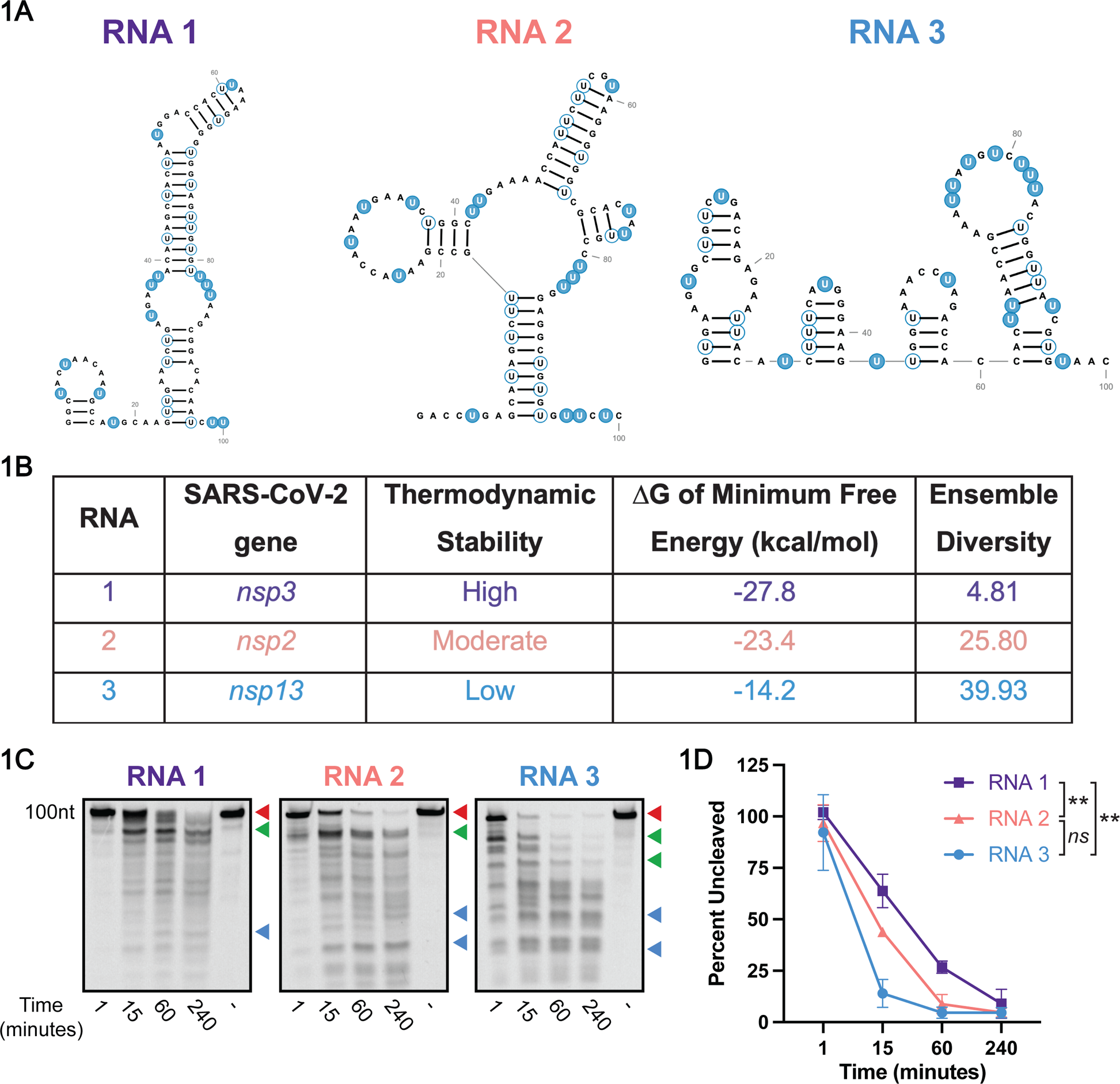
Secondary structure protects RNA from cleavage by SARS-CoV-2 nsp15. **A.** Predicted minimum free energy structures (RNAfold) of 100 nt RNAs from the SARS-CoV-2 genome of varying thermodynamic stability (high, medium, and low). Unpaired uridines are highlighted in blue, and paired uridines are outlined in blue. All RNA structures were predicted in RNAfold and designed in RNA2Drawer (27, 61). **B.** The locations within the SARS-CoV-2 genome are indicated for each structure, and thermodynamic stability is indicated as −ΔG. Ensemble diversity is listed as predicted by RNAfold. **C.** Endonuclease assays of RNAs from (A). Full-length (uncut) RNAs are indicated by the red arrows. Diminishing cleavage products (cleavage intermediates) are indicated by green arrows. Accumulating cleavage products (cleavage end products) indicated by blue arrows. Representative images from one of three total experiments are shown. **D.** Quantitation of data from (C). The percentage of full-length RNA remaining was measured by densitometry. Most thermodynamically stable RNA (RNA 1) was cleaved least rapidly relative to other RNAs. Percentage of uncut RNA was calculated by normalizing to a denatured nsp15 control (indicated by (-) in the far-right lane). Area under the curve (AUC) was calculated for each RNA and one-way ANOVA with multiple comparisons was performed on the AUC. Data represents three independent experiments.

### SARS-CoV-2 nsp15 cleaves unpaired bases in structured RNAs

In vitro biochemical studies by Bhardwaj et al. previously showed that SARS nsp15 specifically cleaves 3’ of unpaired uridines in s2m, a conserved structural element found in the 3’UTR of the SARS genome and some other related CoVs (20, 29, 30). In addition to our results described above (Fig. 1), this study suggests that the context in which the cleaved uridine is presented may be important for determining the cleavage specificity of nsp15. In the case of s2m, the two unpaired uridines (U25 and U30) are located in two unstructured loops (Fig. 2A, highlighted in blue). Structural resolution of s2m has shown that U25, which is located within the GNRNA sequence of the pentaloop of s2m, is highly disordered relative to other nucleotides, and that U30 is also moderately disordered (29). Based on previous findings and the availability of structural information, we posited that s2m would be a tractable model RNA for studying the determinants of cleavage specificity of SARS-CoV-2 nsp15.

**Figure 2.**
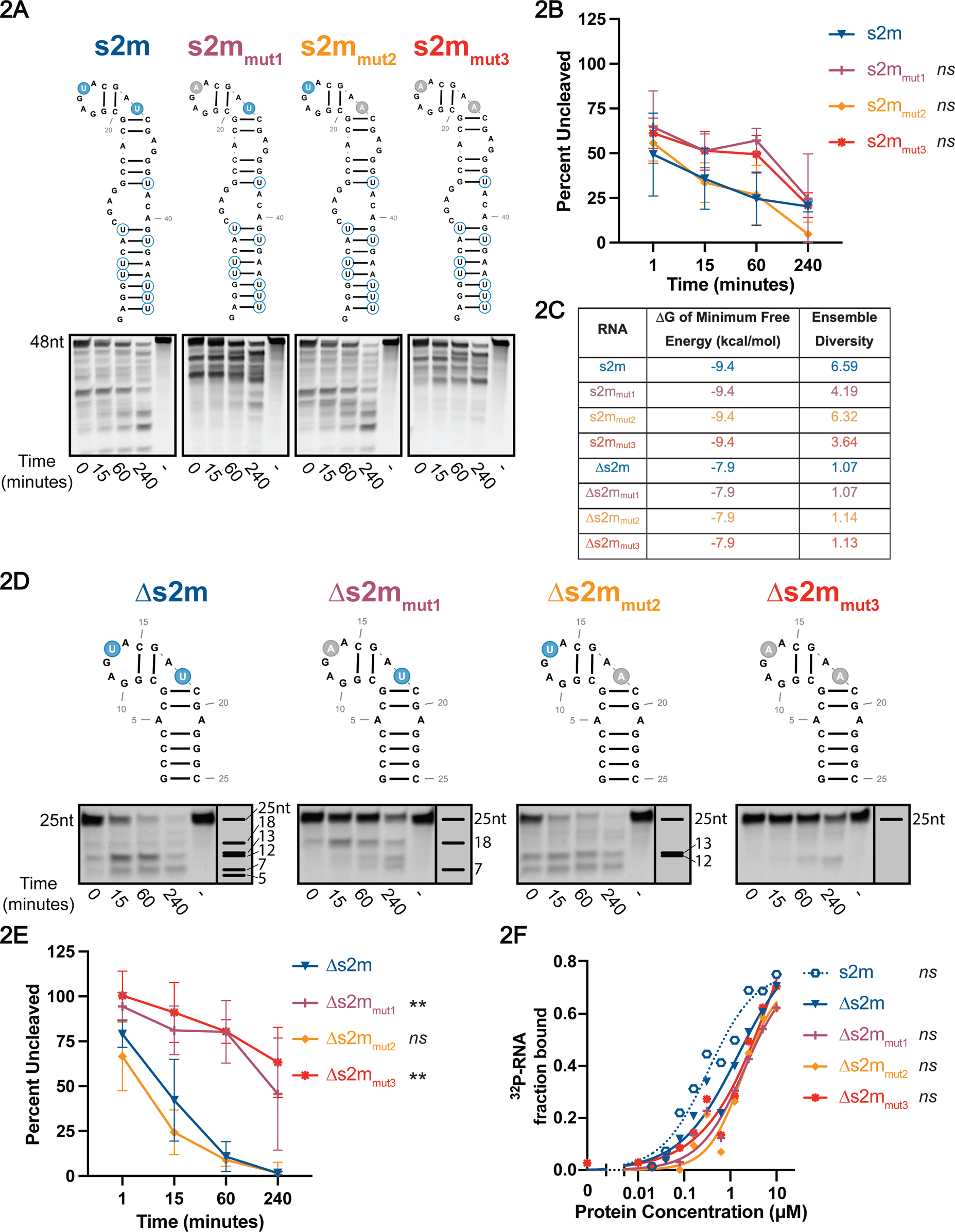
SARS-CoV-2 nsp15 cleaves structured RNAs at specific sites. **A.** Endonuclease assays of full-length s2m mutant RNAs. Full-length bands indicated at the top of the gel with cleavage products below. Representative images from one experiment. **B.** Percent of full-length RNA remaining as measured by densitometry. Percentage of uncut RNA was calculated by normalizing to a denatured nsp15 control (indicated by (-) in the far-right lane). RNAs with uridines present in the pentaloop (s2m and s2m_mut2_) cleaved more efficiently than s2m_mut_1 and s2m_mut3_. AUC was calculated for each RNA and one-way ANOVA with multiple comparisons was performed on the AUC. AUC of s2m mutant RNAs are compared to wt s2m. Data represents three independent experiments. **C.** Thermodynamic stability and ensemble diversity of s2m and mutant RNAs indicated as predicted by RNAfold. **D.** Endonuclease assays of modified s2m RNAs. Full-length bands indicated at the top of the gel with cleavage products below. Representative images from one experiment. **E.** Percent of full-length RNA remaining as measured by densitometry. Percentage of uncut RNA was calculated by normalizing to a denatured nsp15 control (indicated by (-) in the far-right lane). RNAs with uridines present in the pentaloop (Δs2m and Δs2m_mut2_) cleaved more efficiently than Δs2m_mut_1 and Δs2m_mut3_. AUC was calculated for each RNA and one-way ANOVA with multiple comparisons was performed on the AUC. AUC of Δs2m mutant RNAs are compared to wt Δs2m. Data represents three independent experiments. **F.** Nsp15 binding curves of Δs2m RNAs as compared to wt s2m. Δs2m RNA binding curves overlap with each other while wt s2m curve is shifted left indicating improved binding with nsp15. Dissociation constant (K_D_) was calculated for each RNA and one-way ANOVA with multiple comparisons was performed on the K_D_. AUC of wt full-length s2m and Δs2m mutant RNAs are compared to wt Δs2m. Data represents three independent experiments.

We first sought to determine whether SARS-CoV-2 nsp15 exhibits cleavage specificity for unpaired uridines in s2m similar to that previously observed for SARS nsp15 (20). To identify whether nsp15 preferentially cleaves at U25, U30, or both, we generated wild type (wt) SARS s2m RNA, as well as single and double s2m mutants containing U-to-A substitutions at U25 and U30 (Fig. 2A) and compared cleavage efficiency of these RNAs via endonuclease assay (Fig. 2A and B). In contrast to wt s2m, cleavage of the double mutant (s2m_mut3_) was noticeably impaired, consistent with previous studies showing nsp15 cleavage preference for unpaired uridines. However, in contrast to previous findings with SARS nsp15, we observed that mutation of U30 (U30A; s2m_mut2_) did not impact RNA cleavage, whereas mutation of U25 (U25A; s2m_mut1_) did. Although quantitative differences in full-length RNA cleavage were not found to be statistically significant (Fig 2B), accumulation of cleavage products in the U25A (s2m_mut1_) and double mutant (s2m_mut3_) was clearly impaired relative to wt and U30A (s2m_mut2_) RNAs. Interestingly, we also observed accumulation of distinct cleavage products in s2m_mut1_ and s2m_mut3_. This may indicate that step wise cleavage of RNA and RNA intermediates is altered in the absence of the scissile U25. However, while mutation of U25 was not predicted to impact the structure of s2m using RNAfold analysis (Fig. 2C), we cannot rule out the possibility that mutation of U25 leads to changes in s2m RNA structure which impact nsp15-mediated cleavage.

Due to the presence of multiple uridines in s2m, accurate identification of cleavage products and intermediates was challenging. In order to eliminate extraneous cleavage products to clarify interpretation of endonuclease data and further dissect RNA structural determinants of cleavage efficiency, we designed truncated s2m RNAs (Δs2m) which lack the base of the stem and all uridines except U25 and U30 (Fig. 2D). Similar to the experiments described above, we also engineered Δs2m mutants with U-to-A substitutions at U13 (Δs2m_mut1_) and U18 (Δs2m_mut2_) (equivalent to U25 and U30 in wt s2m), or both U13 and U18 (Δs2m_mut3_). As with the full-length s2m RNAs, the free energies and ensemble diversities of the Δs2m mutant RNAs were similar to each other (Fig. 2C). We analyzed cleavage of wt and mutant Δs2m RNAs over time by endonuclease assay as described above (Fig 2D and E). In keeping with experiments with full-length s2m RNA, we observed rapid accumulation of cleavage products in wt Δs2m and Δs2m_mut2_ RNAs. In contrast, mutation of both U13 and U18 (U25 and U30 equivalent), or U13 (U25 equivalent) alone, resulted in delayed and reduced nsp15 cleavage of RNA. Interestingly, we still observed some modest cleavage of Δs2m_mut1_ suggesting that while U13 is the predominant cleavage site in the Δs2m RNA, nsp15 is also able to recognize and cleave at other sites. While mutation of both U13 and U18 led to a further reduction in nsp15-mediated cleavage, some cleavage products were still detected in the double mutant, lending further support to the hypothesis that nsp15 can also cleave non-uridine bases, albeit at lower efficiency. Indeed, while nsp15 displays a strong preference of cleavage 3’ of uridines, previous work has shown that cleavage at other bases (especially cytosine) also occurs (31).

We hypothesized that RNA secondary structure contributes to nsp15 cleavage specificity by presenting the scissile uridine in a structural context favorable for nsp15 cleavage. However, an alternative explanation could be that RNA structure serves to sterically hinder or promote nsp15-RNA binding. To address whether differences we observed in cleavage of wt and mutant Δs2m RNAs was due to differences in protein-RNA binding alone, we compared nsp15-RNA binding affinities using differential radial capillary of action ligand assay (DRaCALA) (32, 33). Here, 5’-end radiolabeled Δs2m RNAs were incubated with increasing concentrations of recombinant mutant nsp15 (nsp15 (K290A)), complexes applied to nitrocellulose membrane, and the fraction of bound vs unbound RNA quantified (Fig. 2F). Nsp15 (K290A) was used to eliminate endonuclease activity and contains a mutation of one of the conserved catalytic residues necessary for endoU activity (20, 34–36). We compared binding of wt and mutant Δs2m RNA, in addition to full-length wt s2m. The latter was chosen as a control, as it is presently unknown what the minimum length requirement is for RNA recognition and binding by nsp15. We observed binding of nsp15 (K290A) to wt Δs2m (K_D_=1.5µM). When we compared binding of K290A to each mutant Δs2m RNA, we observed only slight decreases in binding affinity (approximately 1.3-fold; Δs2m_mut1_ K_D_=2.0µM, Δs2m_mut2_ K_D_=1.9µM, Δs2m_mut3_ K_D_=8.5µM), which were not found to be statistically significant. Notably, we observed a modest increase in binding affinity of K290A for the full-length s2m RNA (approximately 4-fold; s2m K_D_=0.3µM) suggesting that other RNA-protein contacts in longer RNAs may modestly contribute to binding affinity. As nsp15 exhibited similar binding affinities for all Δs2m RNAs we concluded that differences in cleavage efficiency of Δs2m RNAs is not due to differences in protein-RNA binding, but likely driven by positioning of the scissile uridine in these differently structured RNAs.

### Flexible uridine nucleotides in structured pentaloops are susceptible to SARS-CoV-2 nsp15 cleavage

Recent structural studies have suggested that SARS-CoV-2 nsp15 uses a base flipping mechanism to position the scissile uridine in the catalytic pocket, thus enabling cleavage of dsRNA (37). The crystal structure of s2m shows that U25 in the GNRNA pentaloop adopts a similar flipped out orientation (29) suggesting that this mechanism could also account for the specificity of nsp15 cleavage for U25 in s2m. RNA pentaloops adopt a characteristic structure which consists of a sheared G-A base pair that closes the pentaloop and induces base stacking of first N, R, and A nucleotides. Notably, this conformation induces extrusion (or ‘flipping’) of the second N (GNR**<u>N</u>**A) in the pentaloop (38, 39). Thus, we postulated that the position of the scissile uridine in the pentaloop (GAG**<u>U</u>**A) might be necessary to confer the correct orientation required for nsp15 cleavage. To test whether the relative position of U25 is necessary for cleavage specificity, we constructed two additional Δs2m mutants in which the position of scissile uridine has been altered (Fig. 3A, top), and compared cleavage efficiency in the endoU assay (Fig 3A, bottom). In Δs2m_mut4_ the adjacent guanidine and uridine in the pentaloop were swapped (GA**<u>UG</u>**A), such that the scissile uridine now sits at the apex of the pentaloop. In Δs2m_mut5_ the scissile uridine was positioned downstream at the 3’ end of the pentaloop and adjacent to closing GC base pair of the helix (GAGA**<u>U</u>**). The second uridine in the bulge of Δs2m (U18) was also mutated in both RNAs in order to assess the impact of U13 (U25 equivalent in full-length s2m) cleavage only. We predicted that in Δs2m_mut4_ positioning of the scissile uridine at the apex would have minimal effect on cleavage but that placement of uridine adjacent to the helix would place structural constraints on this base such that it would be unable to adopt the necessary conformation (flipped out) required for positioning within the catalytic pocket of nsp15. When we compared cleavage of each mutant, we observed that Δs2m_mut4_ (GA**<u>U</u>**GA) was cleaved to completion earlier than both Δs2m_mut2_ (control) and Δs2m_mut5_ (GAGA**<u>U</u>**) (Figure 3B). In contrast, we observed almost no cleavage of Δs2m_mut5_ supporting the hypothesis that the scissile uridine might be more structurally constrained at this position and unable to engage with the catalytic pocket of nsp15. From these data, we concluded that uridine position within the pentaloop influences nsp15 cleavage activity, likely through structural constraints.

**Figure 3.**
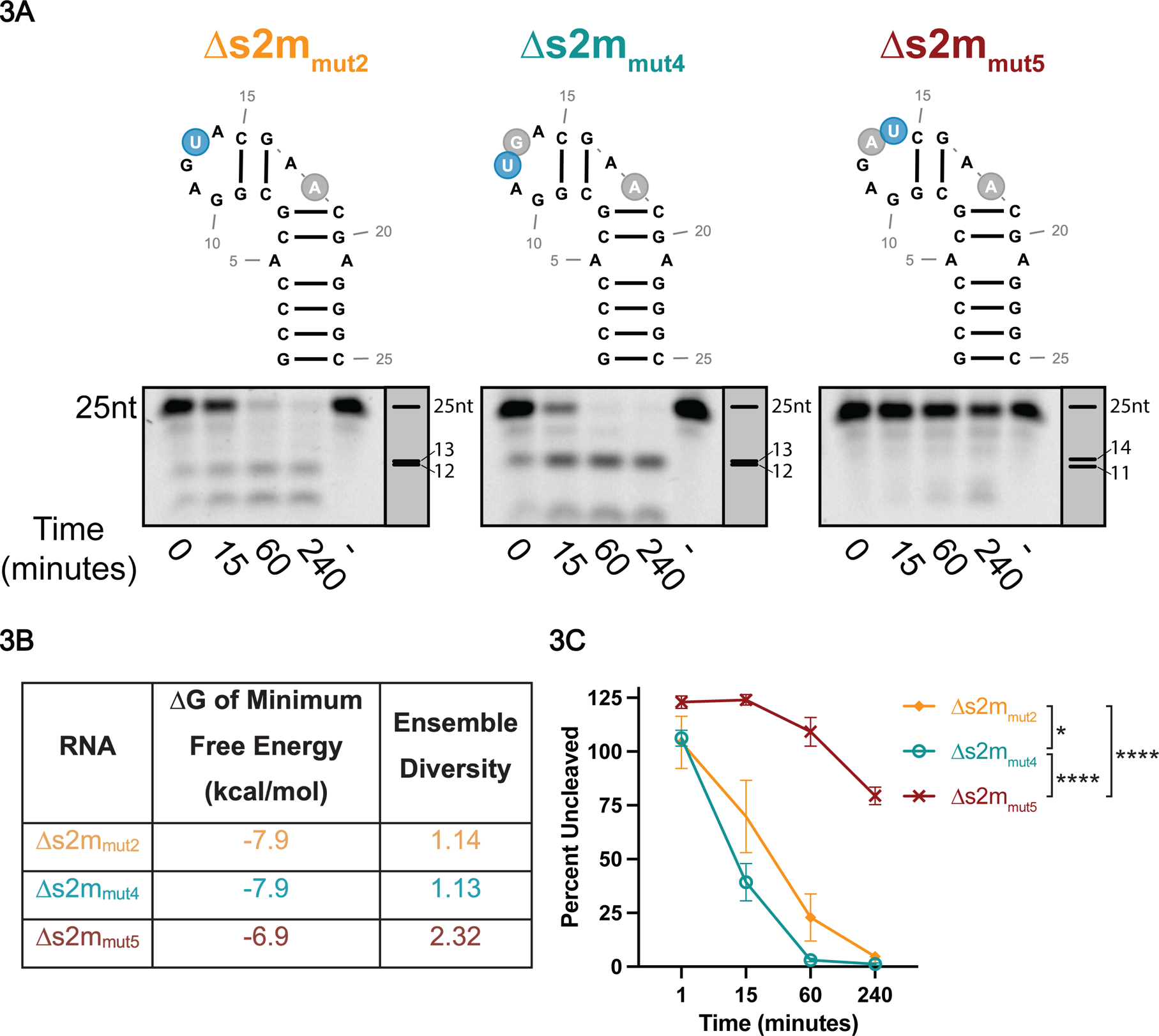
Uracil position in loop structures impacts SARS-CoV-2 nsp15 cleavage efficiency. **A.** Endonuclease assays of s2m pentaloop mutant RNAs. Full-length bands indicated at the top of the gel with cleavage products below. Representative images from one experiment. **B.** Thermodynamic stability and ensemble diversity of s2m and mutant RNAs indicated as predicted by RNAfold. **C.** Percent of full-length RNA remaining as measured by densitometry. Percentage of uncut RNA was calculated by normalizing to a denatured nsp15 control (indicated by (-) in the far-right lane). RNAs with uridines positioned closer to the apex of the pentaloop cleaved more rapidly than Δs2m_mut5_ in which the uridine is at the end of the pentaloop. AUC was calculated for each RNA and one-way ANOVA with multiple comparisons was performed on the AUC. Data represents three independent experiments.

## Discussion

Nsp15 is a viral endonuclease conserved across the *Nidovirales* and plays a crucial role in CoV evasion of host innate immunity. The crystal structure of several CoV nsp15 proteins has been solved to date, including SARS-CoV-2 (34, 36, 40–44). Structural and biochemical analyses have revealed key aspects of nsp15 function, including the requirement of Mn^2+^ for endonuclease activity at neutral pH as well as formation of higher order oligomers (hexamers) which is also necessary for endoU function (20, 34, 35, 40, 44, 45). Nsp15 primarily functions to evade innate immune responses by cleaving dsRNA replicative intermediates and preventing activation of MDA-5, and possibly other RNA sensors in the host cell (16-19, 46-50). Some studies also suggest that CoV nsp15 may play a role in viral transcriptional regulation during viral replication (19) and that endoU-independent activities additionally contribute to modulation of host responses (49, 51–54).

At present, the RNA targets of nsp15 endoU from across the *Coronaviridae* are poorly defined. This can be attributed in part to the fact that in vitro studies have used different RNA species (dsRNA vs ssRNA) and different RNA sequences and lengths, making cross-comparison of these data challenging. Nsp15 has been shown to cleave 3’ of pyrimidines, in particular uridine, and exhibit a preference for weak bases (A or U) 3’ of the scissile nucleotide. However, to date no specific nsp15 recognition motifs have been described. Indeed, our analysis of previously identified MHV nsp15 cleavage sites (19) failed to identify novel sequence motifs in and around the cleavage site (data not shown). In the absence of a sequence motif, we postulate that RNA secondary structure may function to regulate both RNA binding and cleavage efficiency of nsp15.

To this end, recent work has shown that SARS-CoV-2 nsp15 manipulates the scissile uridine for cleavage of dsRNA (37). However, the extent to which RNA secondary structure is involved in modulating SARS-CoV-2 nsp15 cleavage activity remains to be determined. Thus, in this study we sought to determine the role of RNA structure in the regulation of SARS-CoV-2 nsp15-mediated cleavage and define specific structural features of RNAs that contribute to this. We demonstrated that increased thermodynamic stability of RNA is associated with decreased nsp15 cleavage efficiency (Fig. 1) and postulated that stable secondary structure may contribute to nsp15 cleavage by two modes: 1) steric hinderance; and 2) inducing optimal positioning of the scissile uridine in the nsp15-RNA complex. To test this second hypothesis, we used s2m as a model RNA to explore the RNA determinants of nsp15 cleavage specificity.

Using a genetic approach, we first demonstrated that nsp15 exhibits specificity for cleavage of the uridine nucleotide located in the GNRNA pentaloop of s2m (Fig. 2, Fig. 3). This contrasts with previous data from Bhardwaj et al., in which the authors observed preferential cleavage of U30, located in the bulge sequence downstream of the pentaloop (20). It should be noted that these previous studies were performed with SARS nsp15, which shares 88% identity with SARS-CoV-2 nsp15. One explanation for the discrepancy in our data could be structural differences between these proteins which lead to altered cleavage specificity.

Notably, the specificity for cleavage of the pentaloop uridine (GAG**<u>U</u>**A) was not due to differences in nsp15-RNA binding (Fig. 2F), thus we postulated that the sequence context of the scissile uridine might be important for this specificity. The characteristic structure adopted by GNRNA pentaloops induces extrusion, or base flipping, of the scissile uridine away from the body of the RNA (38, 39), and this base flipping mechanism has been suggested to be important for correct positioning of uridine in the catalytic pocket of nsp15 (55). Pentaloop motifs have also been implicated in base editing mediated by <u>A</u>denosine <u>D</u>eaminases that <u>A</u>ct on <u>R</u>NA 2 (ADAR2) (56), though in this instance the GCUMA pentaloop is located downstream of the edited A, and is implicated in recruitment of ADAR2 via interactions between the dsRNA binding motif of ADAR2 and the target RNA. We speculated that if structural conformation of the scissile uridine was important for base recognition and cleavage that altering the position (and conformation) of the uridine base would lead to altered cleavage efficiency. Indeed, repositioning of the scissile uridine to the apex of the pentaloop (Δs2m_mut4_) slightly enhanced cleavage efficiency RNA, whereas repositioning adjacent to the closing G-C base pair (Δs2m_mut5_) significantly reduced RNA cleavage (Fig. 3). The marked decrease in Δs2m_mut5_ cleavage is likely due to altered positioning of the scissile uridine which we predict is no longer extruded from the pentaloop, as well as decreased flexibility of this base which is now structurally constrained by the neighboring G-C base pair. Another possibility is that cleavage efficiency is impacted by altering the sequence context of the scissile uridine. Nsp15 has been shown to preferentially cleave uridines flanked by a weak base (A/U) which is found in wt Δs2m and Δs2m_mut2_ (U^↓^A) but is changed to a strong base in Δs2m_mut5_ (U^↓^C). However, we observed a modest increase in cleavage efficiency of Δs2m_mut4_ which also has a strong base at the 3’ position (U^↓^G) and lends support to the hypothesis that the structural context of the scissile uridine, not sequence context, may be more relevant in dictating cleavage specificity and efficiency.

Collectively, our data suggests that RNA secondary structure plays two distinct roles in determining nsp15 cleavage specificity. First, thermodynamically stable structures likely sterically hinder nsp15-RNA interactions, thus preventing engagement and cleavage of some RNA sequences. Second, susceptible bases are presented in a structural context which facilitates optimal positioning of the scissile nucleotide in the catalytic pocket of nsp15 (e.g., extrusion of the scissile U25 in the GNRNA pentaloop). However, several questions remain unanswered, most notably, what are the biologically relevant targets (host and viral) of nsp15 cleavage during CoV infection and are viral targets conserved across different CoVs? Two studies to date have identified cleavage sites within the positive-sense genomic RNA and 3’ polyU tails of the negative-sense RNA of MHV (17, 19). Despite this, no recognition motif or other regulatory element associated with nsp15 cleavage has been identified. While the in vitro assays performed here and in other studies provide a tractable system for dissecting the molecular mechanism of nsp15-RNA interactions and cleavage determinants, nsp15 forms part of a larger replication-transcription complex (RTC) which contains other viral proteins and host factors. How nsp15 protein-protein interactions and RNA interactions with other replicase proteins (e.g., nsp12, nsp14, nsp16) impacts nsp15 cleavage of RNA still remains unsolved. Indeed, recent models of the RTC suggest the hexameric structure of nsp15 provides the central structure and arrangement of a hexameric RTC (57). When nsp15 is bound in the RTC structure, there may be differences in its cleavage of viral RNAs due to its 3-D interaction with the nascent RNA as opposed to when nsp15 is free from the RTC as was the case in this study. The importance of examining nsp15-RNA interactions in their relevant biological context is further highlighted by recent studies showing that the requirement for Mn^2+^ and the structural conformation of nsp15 is highly dependent on the pH of the environment (58). Given that the redox state of the viral RTC of +ssRNA viruses is altered during replication, the conditions under which nsp15 endoU activity is examined is highly relevant. To address this our future studies will explore the impact of pH and redox state on nsp15 structure and function in vitro.

Overall, our findings suggest a role for RNA secondary structure in regulation of nsp15 cleavage. Identification of bona fide nsp15 cleavage sites in SARS-CoV-2 RNA during replication and examination of how perturbation of these sites impacts the outcome of replication and pathogenesis will provide key insight into the broader roles for nsp15 endoU activity during infection (e.g., viral transcriptional/translational regulation, host translation). Furthermore, defining key structural features of RNAs which are resistant or susceptible to nsp15 cleavage may provide avenues for development of antiviral therapeutics.

## Materials and Methods

*Generation of SARS-CoV-2 RNA segments:* 100nt segments of SARS-CoV-2 genome were amplified using the primers listed in Table 1. Amplicons were purified using Mag-Bind Total Pure NGS beads (Omega Bio-Tek). 1.5 x bead volume relative to amplification reactions was used and purified as per manufacturer’s instructions. Purified amplicons were subcloned into a vector containing the HDV ribozyme (59). These constructs were linearized and purified prior to transcription.

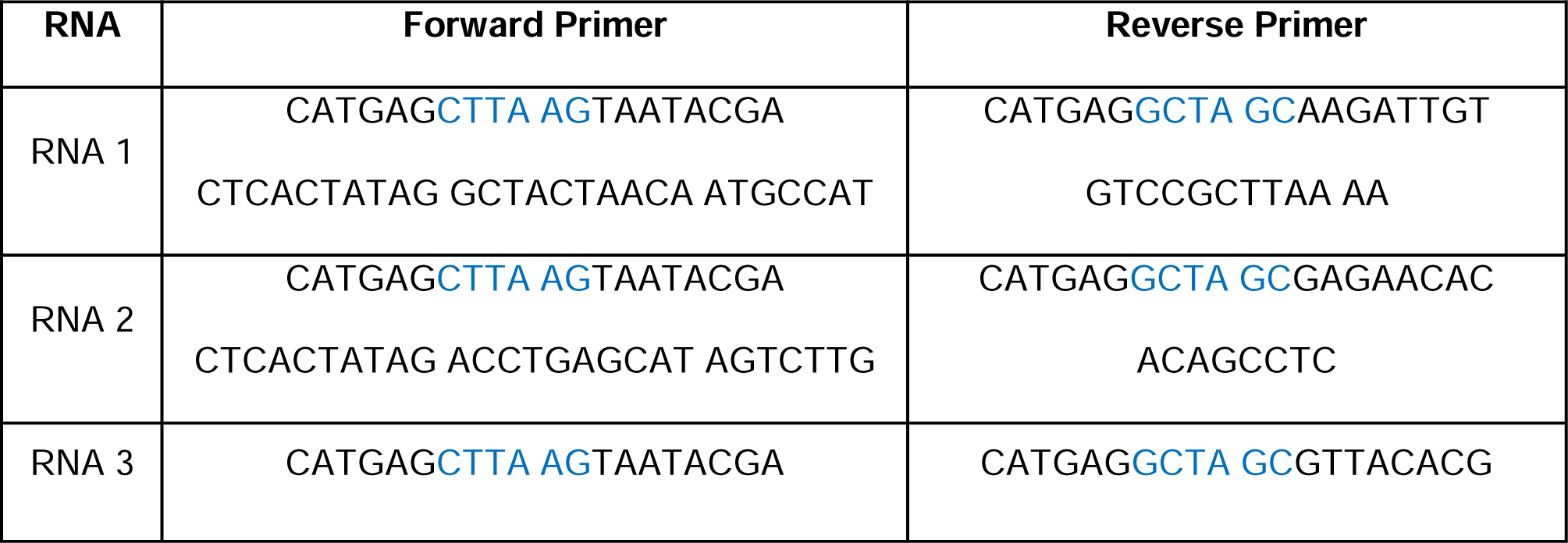
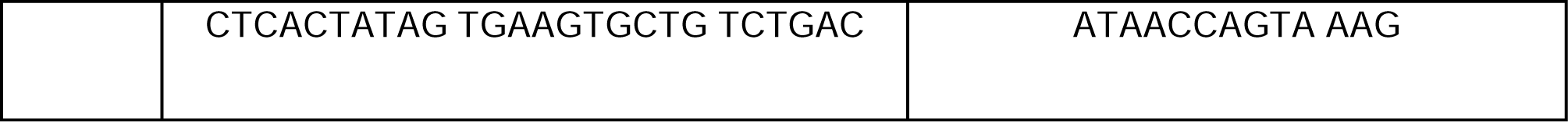
**Table 1**

Approximately 5 pmol of purified DNA was used as a template for T7 transcription reactions. RNAs were transcribed using the HiScribe T7 High Yield RNA Synthesis Kit (New England Biolabs). Reactions were set up as per manufacturer’s instructions with the addition of 4 units of Murine RNase Inhibitor (NEB) and 5% DMSO. Transcriptions were incubated at 42°C overnight after which, the reactions were treated with DNaseI (NEB) as per manufacturer’s instructions. DNase-treated transcriptions were then purified using Mag-Bind Total Pure NGS beads as described above.

Transcriptions were gel purified by running samples on urea gels. 15% TBE-Urea gels were made using SequaGel UreaGel 29:1 Denaturing Gel System (National Diagnostics) Urea was added to the samples to a final concentration of 4M before running on the gels at 200V for 90 minutes. Gels were briefly stained with 0.02% Methylene Blue in TBE until ladder was visible. Gels were imaged on Gel Doc XR+(Bio-Rad), and bands corresponding to 100nt fragments were cut out and shredded by centrifugation. RNAs were eluted overnight on rotator in 3x volume of elution buffer (10mM Tris-HCl, 1mM EDTA, 300mM NaCl, 0.1% SDS) relative to gel weight. Gel slurries were applied to cellulose acetate columns and frozen at −80°C for 10 minutes. Samples were centrifuged at 10,000x g for 3 minutes. Eluate was adjusted up to 600µL with nuclease-free water and purified by acid phenol-chloroform extraction.

[utbl1]

### Endonuclease Assay

Indicated amounts of RNAs were diluted in folding buffer (50mM Tris-HCl, 50mM KCl, 5mM MgCl_2_, 5mM DTT). RNAs were incubated at 95°C for 10 minutes then cooled to 25°C at a rate of 0.1°C/sec. Purified nsp15 was diluted in assay buffer (50mM Tris-HCl, 50mM KCl, 50mM MnCl_2_, 5mM MgCl_2_, 5mM DTT) to the appropriate concentration for indicated nsp15:RNA ratios. Recombinant nsp15 was expressed and purified as previously described (60). To denature nsp15, the protein was incubated at 95°C for 5 minutes. Folded RNA was aliquoted into individual tubes (1 tube/timepoint), and diluted nsp15 was mixed in equal amounts to each RNA simultaneously. Reactions were arrested at indicated timepoints with a final concentration of 20mM EDTA before flash freezing in liquid nitrogen. Denatured nsp15 reactions were incubated for 240 minutes.

TBE-Urea gels were made using SequaGel UreaGel 29:1 Denaturing Gel System (National Diagnostics). 15% or 22.5% gels were poured as indicated in each experiment. RNA samples were thawed and mixed with an equal volume of 2x RNA Loading Dye (NEB) and loaded onto the gels. Gels were run at 200V for 1 hour in 1x TBE. Gels were stained with SYBR Green (ThermoFisher) diluted 1:5000 in TBE for 30 minutes while shaking and imaged on Gel Doc XR+(Bio-Rad). Full-length bands for at each time point were calculated by densitometry using Image Lab software (Bio-Rad). Density of bands were normalized to denatured nsp15 control for each reaction.

### DRaCALA

DRaCALAs were performed as described previously (32). Briefly, RNAs were dephosphorylated and 5’ radiolabeled with 30µCi ATP. Radiolabeled RNAs were set up in reactions with the following components: DRaCALA buffer (50mM HEPES, 100mM KCl), 5mM MgCl_2_, 50ng/µL yeast tRNA, 1mM DTT, 5% glycerol. RNAs were heated to 95°C for 2 minutes and cooled at RT for 10 minutes. Serial dilutions of nsp15 K290A were added to the reactions which were incubated for 15 minutes at RT and spotted onto a 0.45µm nitrocellulose membrane. Blots were exposed to a phosphor screen overnight and imaged on Sapphire Molecular Imager (Azure Biosystems). Binding was measured by densitometry as previously described (33) using Fiji.

## Acknowledgements

The authors declare that they have no conflict of interest. This project has been funded in part with Federal funds from the National Institute of Allergy and Infectious Diseases, National Institutes of Health, Department of Health and Human Services, under Contract No.: 75N93022C00036.

